# Integration of hyperspectral imaging and transcriptomics from individual cells with HyperSeq

**DOI:** 10.1101/2024.01.27.577536

**Authors:** Yike Xie, Abbas Habibalahi, Ayad G. Anwer, Kanu Wahi, Catherine Gatt, Emma M. V. Johansson, Jeff Holst, Ewa Goldys, Fabio Zanini

**Affiliations:** School of Clinical Medicine, UNSW Sydney, Australia; School of Biomedical Engineering, UNSW Sydney, Australia; School of Biomedical Sciences, UNSW Sydney, Australia; Flow Cytometry Unit, Mark Wainwright Analytical Centre, UNSW Sydney, Australia; UNSW Cellular Genomics Futures Institute, Sydney, Australia; Evolution & Ecology Research Centre, School of Biological, Earth & Environmental Sciences, UNSW Sydney, Australia

**Author notes:** Equal contribution (Ewa Goldys, Fabio Zanini).

## Abstract

Microscopy and omics are complementary approaches to probe the molecular state of cells in health and disease, combining granularity with scalability. While important advances have been achieved over the last decade in each area, integrating both imaging- and sequencing-based assays on the same cell has proven challenging. In this study, a new approach called HyperSeq that combines hyperspectral autofluorescence imaging with transcriptomics on the same cell is demonstrated. HyperSeq was applied to Michigan Cancer Foundation 7 (MCF-7) breast cancer cells and identified a subpopulation of cells exhibiting bright autofluorescence rings at the plasma membrane in optical channel 13 (*λ*_ex_ = 431 nm, *λ*_em_ = 594 nm). Correlating the presence of a ring with the gene expression in the same cell indicated that ringed cells are more likely to express hallmark genes of apoptosis and less likely to express genes associated with ATP production. Further, correlation of cell morphology with gene expression suggested that multiple members of the spliceosome were downregulated in larger MCF-7 cells. Multiple genes were evenly expressed across cell sizes but also exhibited higher usage of specific exons in larger or smaller cells. Finally, correlation between gene expression and fluorescence within the spectral range of Nicotinamide adenine dinucleotide hydrogen (NADH) provided insight into the metabolic states of MCF-7 cells. These observations provided a link between the cell’s optical spectrum and its internal molecular state, demonstrating the utility of HyperSeq to study cell biology at single cell resolution by integrating spectral, morphological and transcriptomic analyses into a single, streamlined workflow.

## Introduction

Single cell omics are accelerating our understanding of cell biology^1^, shedding light on new cell types and states in culture as well as *in vivo* across tissues, organs, and organisms ^2–5^. While major breakthroughs have been achieved on increasing throughput of scRNA-Seq and related technologies to thousands ^6–9^ and sometimes millions ^9,10^ of cells and to integrate distinct omic modalities into a single experiment (e.g. CITE-Seq ^11^, scATAC+scRNA-Seq ^12^), progress in combining omics with additional, complementary measurements from the same cell has been more limited ^13–16^. Multimodal characterisation of individual cells would benefit biomedical applications including cancer and infectious disease, given that both malignant and virus-infected cells can be genetically ^17^, transcriptionally ^18^, and metabolically ^19^ diverse within a single culture.

Fluorescence-based microscopy has been used to explore cellular heterogeneity for decades. Epifluorescence instruments are commonly used to detect the incoherent light emitted by extrinsic macromolecules including genetically encoded alloproteins (e.g. green fluorescent protein) and artificial dyes (e.g. DAPI). Hyperspectral imaging, which uses narrow band optical filters on both excitation and emission frequencies to detect the autofluorescence of intrinsic biomolecules, has emerged in recent years as a particularly information-rich methodology to study cytometabolic heterogeneity ^20–22^. Hyperspectral imaging can be used to assign to each cell multiple features related to morphology, spectral profile, and combinations of the two (e.g. spectral features with a specific subcellular localization). Hyperspectral data can also be used to train machine learning models aimed at identifying measurable optical proxies of biomedically relevant phenotypes, e.g. to distinguish cancerous cells from normal ones ^23,24^. Nonetheless, connecting hyperspectral features with defined and interpretable molecular states such as apoptosis, activation, or cell cycle remains challenging.

Multiple groups have attempted to combine optical and sequencing methods at the single-cell level into a single experimental approach ^25,26^. Spatial transcriptomic technologies have advanced tremendously in recent years, with commercial solutions such as Visium and Xenium (10X Genomics), Slide-Seq ^27^, and STOmics (BGI). These methods aim to capture large areas of *ex vivo* tissues. This requirement can only be met by today’s technologies by accepting significant tradeoffs either on spatial resolution or on molecular sensitivity.

Techniques to achieve both high transcriptome quality and high spatial resolution have been proposed. Live-seq uses an atomic force microscope with a hollow cantilever tip to extract only a fraction of the cytoplasmic RNA from a living cell ^13^. Its platform, FluidFM ^28^, requires a high level of expertise and currently offers limited throughput in terms of cell numbers. Laser dissection and capture suffer from similar limitations in terms of throughput and user skill requirements ^29,30^. Microfluidic droplet generators such as Drop-Seq ^6^, inDrop ^31^ and commercial solutions (e.g. 10X Genomics, BD Rhapsody) can be coupled to a microscope ^32^, however the droplets are not addressable, therefore the link between cell image and transcriptome is lost. Fluidigm C1 microfluidic chips ^33^ and the Takara ICELL8 ^34^ system can be used in theory to relate optical and transcriptomic features from the same cell. However, both technologies have limited image quality and are used primarily to verify cell capture.

Here, we report the development of HyperSeq, a novel approach that combines hyperspectral autofluorescence imaging with transcriptomics from the same cell and scales to hundreds of cells in a single experiment. We applied HyperSeq to Michigan Cancer Foundation 7 (MCF-7) cells, the most widely studied human breast cancer cell line ^35^, and correlated both spectral and morphological features of each cell with gene expression and preferential exon usage. We discovered that cells with spectrally distinctive bright rings located at the plasma membrane expressed higher levels of mitochondrial transcripts and of nuclear transcripts involved in gene silencing and apoptosis. We then focused on non-apoptotic cells and compared cell morphology with gene expression. We found that cell size is correlated with differential expression of multiple spliceosome components and with preference for specific exons within the same gene. Finally, we found that fluorescence intensity around the peak of Nicotinamide adenine dinucleotide hydrogen (NADH) can suggest metabolic conditions within the cell. HyperSeq has lower technical requirements, better image quality, and higher throughput than other methods, opening new avenues in the identification of cell states and behaviour.

## Results

### Designing a workflow for single cell imaging and transcriptomics

To characterise both optical and transcriptomic features within the same cell, a workflow called HyperSeq, which integrates microscopy and scRNA-Seq, was designed (**Figure 1A**, see Methods for a detailed protocol). Briefly, breast cancer MCF-7 cells ^35^ were seeded onto a gridded cell culture dish (ibidi, Gräfelfing, Germany) at low density and imaged on an Olympus IX83 inverted microscope customised for collection of brightfield and 15-colour hyperspectral autofluorescence (**Supplementary Table 1**, see **Figure 1A** for a representative false-colour image of channel 2). After imaging, each cell culture was transferred to a different laboratory, where individual cells were isolated using an automated cell picker (CellCelector, Sartorius AG, Germany) and deposited into a 384-well PCR plate. Low-resolution brightfield images before and after capture were used to verify successful cell isolation (**Figure 1A**). Smart-seq2 was scaled down for cost effective transcriptome library preparation ^18^. A pilot experiment using 30 cells with and without imaging steps indicated that imaging and picking had a negligible impact on RNA quality as evaluated by mapped read counts (**Supplementary Figure 1A**) and fraction of mitochondrial to total reads (**Supplementary Figure 1B**).

**Figure 1:**
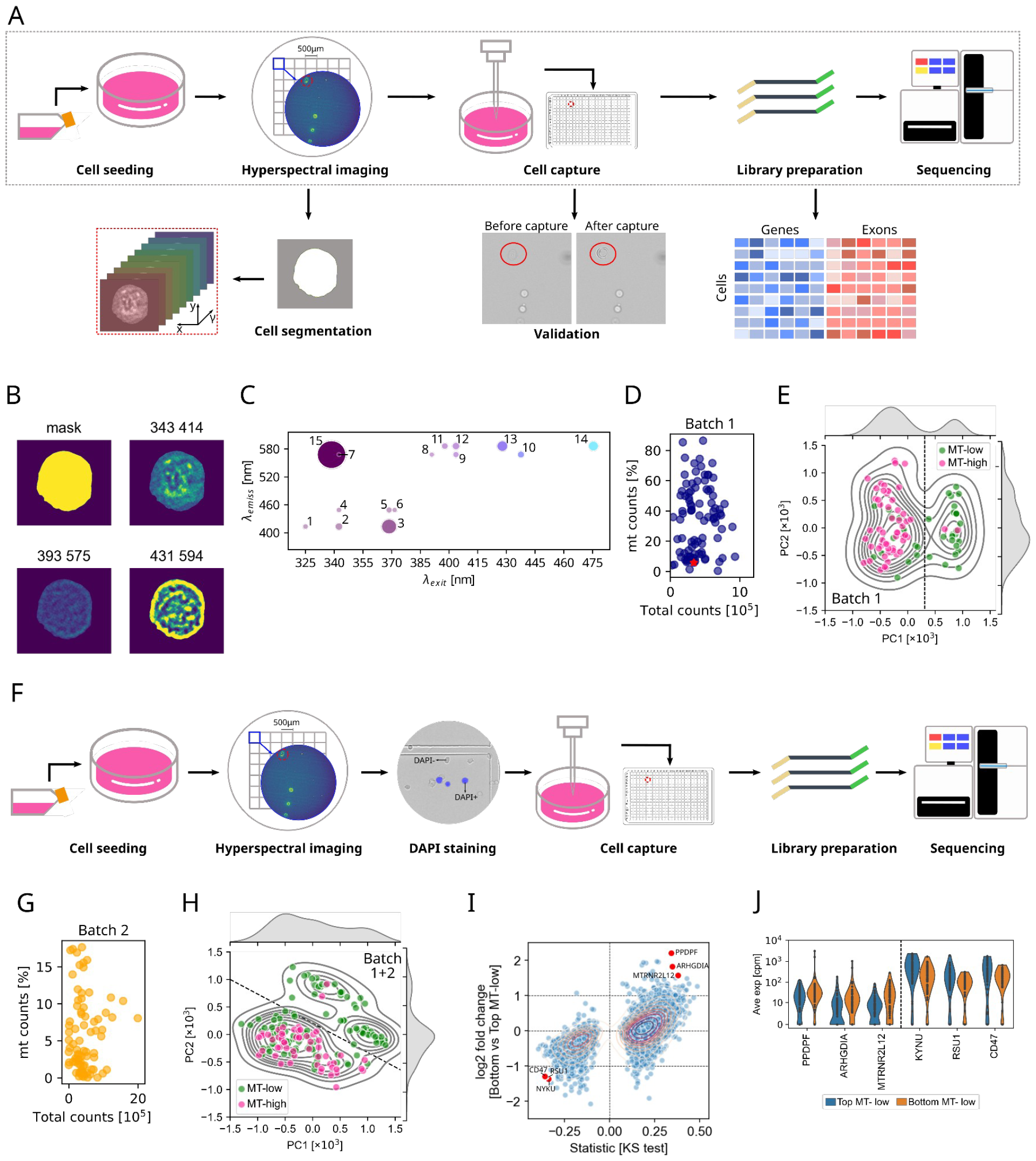
HyperSeq integrates hyperspectral imaging and transcriptomics from the same cell. **A**, Schematic of HyperSeq’s experimental and computational workflow. **B**, Segmentation mask and select autofluorescence channels for a representative cell. **C**, Autofluorescence spectrum for the same cell as in (B). Each dot indicates a channel with the channel number labelled (**Supplementary Table 1**). Dot size indicates fluorescence intensity detected in each channel and dot colour is a left-right gradient corresponding to excitation wavelength. All pixels within the segmentation mask were averaged. **D**, Total reads per cell versus percentage of mitochondrial over total reads for experimental batch 1, with the same cell as in (B) highlighted as a red star. **E**, Hyperspectral principal component analysis (PCA) of experimental batch 1 highlighting two groups of cells, divided by a straight line. **F**, Improved experimental workflow with a DAPI imaging step before picking to identify low-viability cells. **G**, As in (D) but for experimental batch 2. **H**, Hyperspectral PCA for both batches combined, highlighting two groups of cells (upper-right and lower-left). **I**, Scatter plot of the log2 fold change between MT-low cells in lower-left and upper-right subgroups (y-axis) and signed statistics from ks test (x-axis). Genes that are expressed with less than 10 cells are filtered out. Some genes that are lower or higher expressed in bottom MT-low cells are labelled. **J**, Violinplot showing the expression of highlighted genes in (I) in top and bottom MT-low cells.

### HyperSeq integrates optical and molecular information at the single cell level

Given the promising outcome of the initial feasibility study, a larger batch of 126 MCF-7 cells was collected. To quantify the hyperspectral images, a semi-automated cell segmentation pipeline using fake colours was developed (see Methods and **Supplementary Figure 1C**). The segmentation mask was confirmed visually and produced images with diverse patterns at subcellular resolution (**Figure 1B**). To quantify the spectral information of each cell, the optical intensity from all pixels within the segmentation mask was averaged and combined into a cell-specific spectral fingerprint, or cell spectrum (**Figure 1C)**.

Quality controls were implemented to verify the efficiency of HyperSeq, including visual confirmation of a single cell in the brightfield imaging channel (see Methods). Out of 126 cells, 3 lacked good hyperspectral data and 30 were determined to be doublets upon further visual inspection. Of the remaining 93 single cells that passed quality controls, 55 cells (59%) contained more than 25% of mitochondrial over total reads, an indicator of suboptimal cell viability ^36^ (**Figure 1D**). To further examine this mitochondrial heterogeneity, the hyperspectral images were analysed using principal component analysis (PCA) ^37^, a form of unsupervised learning (**Supplementary Table 1**). When represented on the first two hyperspectral principal components, two groups of cells were observed (**Figure 1E**, separated by a straight line), confirming the suspicion that a subpopulation of cells might exhibit low viability due to the experimental protocol. To test this hypothesis, we added one step to the HyperSeq workflow in which cell viability is tested before capture using DAPI, a marker of cell membrane breaching and death (**Figure 1F**). This step does not affect the hyperspectral images and makes it possible to pick only DAPI-negative cells. A second experimental batch on 83 cells included this step (**Supplementary Figure 1D**). Out of 81 cells passing quality controls, all had less than 25% mitochondrial over total reads (**Figure 1G**).

Hyperspectral PCA on both experimental batches combined again showed two groups of cells. MT-low cells (<25% mitochondrial over total reads) populated both groups (**Figure 1H**, groups separated by a straight line). Within MT-low cells, the lower-left group was compared with the upper-right group for number of detected genes and percentage of ribosomal reads via two-tailed Kolmogorov-Smirnov (KS) tests (**Supplementary Figure 1F**). MT-low cells in the lower-left group showed a higher number of genes than MT-low cells in the upper-right group, while the percentage of ribosomal reads was similar. Differential expression analysis between the two groups of MT-low cells was also conducted (**Figure 1I**).The top three upregulated genes in lower-left vs upper-right MT-low cells were PPDPF, involved in cell differentiation, Rho GDP Dissociation Inhibitor ARHGDIA, and MTRNR2L12, involved in negative regulation of the execution phase of apoptosis (**Figure 1I-J**). Downregulated genes in lower-left vs upper-right MT-low cells include kynureninase (KYNU), RSU1, which encodes a Ras Suppressor protein, and CD47, which encodes an adhesive protein mediating cell-to-cell interactions (**Figure 1I-J**).

Overall, the combination of hyperspectral and transcriptomic information, with the added DAPI quality control step to remove apoptotic cells, enabled the detection of a population of cells that may be pre-apoptotic. This additional step led to an improved protocol with comparable throughput and flexibility.

### Characterisation of cells with hyperspectral rings at the plasma membrane

Because HyperSeq has subcellular optical resolution, we asked if the principal components of each cell’s spectrum are related to spatial heterogeneity within each cell. The explained variance exhibited a gap after principal component 2 (PC2) (**Supplementary Figure 2A**), therefore we focused on the first two components as suggested by random matrix theory ^38,39^. Interestingly, PC1 and PC2 lent themselves to an intuitive optical interpretation: blue-emitting channels had larger squared loadings for PC1 and red-emitting channels for PC2 (**Figure 2A**). Therefore, we examined their hyperspectral images further in channels 3 (*λ*_ex_ = 370nm, *λ*_em_ = 414nm) and 13 (*λ*_ex_ = 431 nm, *λ*_em_ = 594 nm), which had the largest squared loadings onto PC1 and PC2 (**Figure 2B**). Some cells exhibited a ring at the plasma membrane in channel 13 (**Figure 2B**, lower panel). A computational algorithm was built to classify cells with or without bright rings (see Methods and **Supplementary Figure 2B**). It categorised 60 cells (34%) as ringed and 114 cells (65%) as unringed, which was confirmed visually. The majority of ringed cells (85%) were located in the lower-left group of the hyperspectral PCA (**Figure 2C**).

**Figure 2:**
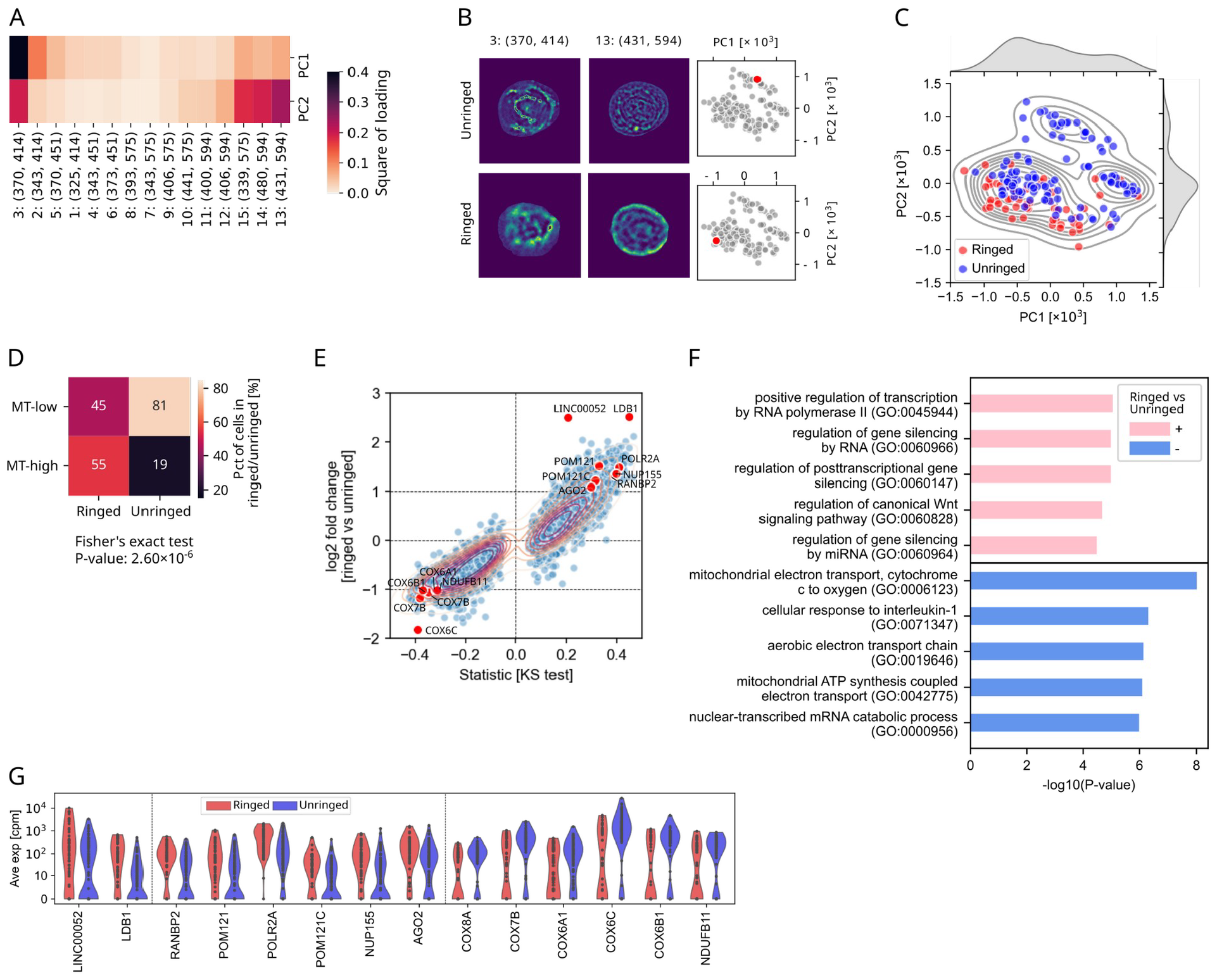
Optical cell heterogeneity can be used as a guide to query transcriptional heterogeneity. **A**, The square of loading of every channel to PC1 and PC2 generated from PCA on average autofluorescent intensities under 15 channels of all cells. **B**, Examples of segmented cells without (upper panel) and with (lower panel) bright rings in channel 13 (*λ*_ex_=431 nm, *λ*_em_=594 nm). The location of example cells in the PC1-PC2 map are shown in two right panels. **C**, Scatter and KDE plots of PC1 and PC2 of each cell. Red and blue face colours indicate cells with (ringed) and without (unringed) rings. **D**, Heatmap showing the percentage of ringed and unringed cells that are MT-low or MT-high. **E**, Scatter plot of the log2 fold change between ringed and unringed cells (y-axis) and signed statistics from ks test (x-axis). Genes that are expressed with less than 10 cells are filtered out. **F**, The top 5 gene ontology (GO) terms among genes upregulated or downregulated in ringed cells relative to unringed cells. Upregulated genes indicate genes with log2 fold change > 1 and -log_10_(p-value)>2, while downregulated genes indicate genes with log2 fold change < -1 and -log_10_(p-value)>2. Genes with median number in either subgroup as 0 were filtered out, and p-values were corrected by the Benjamini/Hochberg (non-negative) method. **G**, Violinplot showing the expression of some genes up- or down-regulated in ringed cells.

Moreover, the majority of ringed cells were MT-high, unlike unringed cells (**Figure 2D**, Fisher’s exact test P-value 2×10^−6^). Direct comparison between ringed and unringed cells revealed differentially expressed genes with a large fold change and a strong statistical support (KS test, **Figure 2E**). Enriched pathways among upregulated genes in ringed (+) and unringed (-) cells were computed via GSEApy ^40^ (**Figure 2F**). Three pathways enriched among genes upregulated in unringed cells were associated with mitochondrial electron transport, indicating a more active tricarboxylic acid (TCA) cycle and increased ATP production ^41^. Genes within these pathways included COX8A, COX7B, COX6A1, COX6C, and COX6B1 encoding cytochrome C oxidase subunits, along with NDUFB11, which encodes ubiquinone oxidoreductase subunit B11 (**Figures 2F-G**). Genes upregulated in ringed cells had a less direct interpretation and were related in part to gene silencing (**Figures 2E-G**). These data indicated that unringed cells might be metabolically more active, whereas ringed cells might have been in a less vital state at the time of sampling. More generally, HyperSeq was effective at discovering subcellular optical features and generating data-driven hypotheses about their biological meaning.

### Cell size correlates with microtubule growth and RNA splicing and degradation

HyperSeq can be used not only to investigate autofluorescence, but also to associate optical features in the brightfield channel, such as cell size and morphology, with gene expression. MCF-7 cells picked from the same culture dish were heterogeneous in size. The largest cell (15,61um^2^) had an area nine times larger than the smallest one (1,66um^2^) (**Figure 3A, Supplementary Figure 3A**). In a cycling cell line, pre-mitotic cells are expected to be roughly twice as large as cells after division. Therefore, additional sources of heterogeneity beyond cell cycle might exist in the culture and be associated with detectable gene expression signatures.

**Figure 3:**
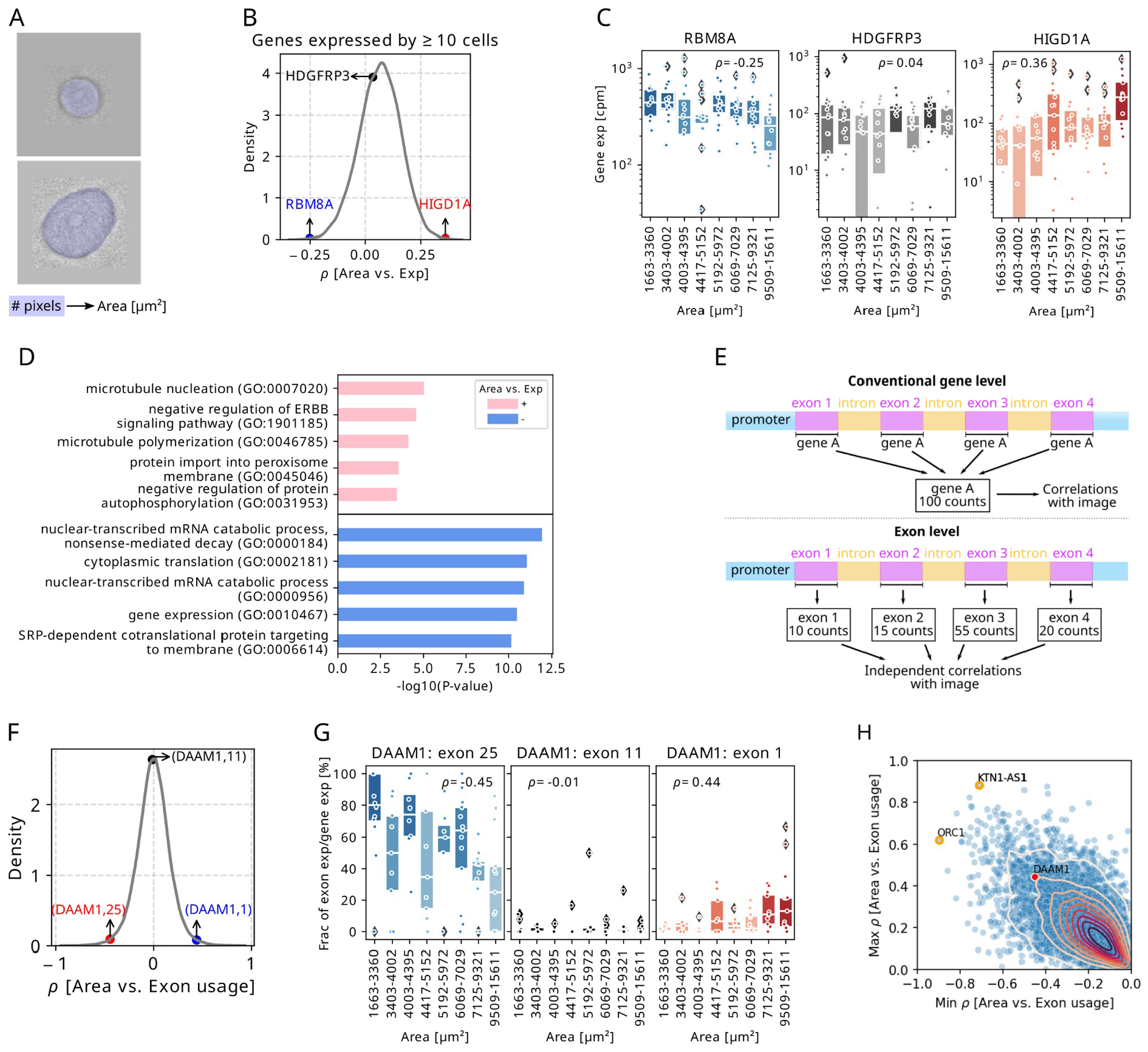
Correlating cell size with gene expression and relative exon usage. **A**, Schematic plot of computing cell areas based on the number of pixels. **B**, Kernel density estimate (KDE) plot of correlation coefficients between cell area and gene expression for genes expressed by ≥ 10 cells. Three genes with negative, ∼ zero, and positive correlation coefficients are labelled. **C**, Box plots showing the gene expression (cpm) in every 15 cells ranking with cell areas from smallest to largest of genes in (**B**). Each dot represents a cell, and box colour is coded by the median expression in each bin. Box plots’ horizontal lines indicate the first, second (median) and third quartiles. **D**, The top 5 gene ontology (GO) terms among 300 genes with highest positive or negative correlation coefficients with cell areas. Genes that are expressed with less than 10 cells are filtered out. **E**, Schematic plot of transcriptomic analysis at gene level and exon level. **F**, KDE plot of correlation coefficients between cell area and exon usage on exons from genes expressed by ≥ 10 cells. Three exons within one gene with negative, ∼ zero, and positive correlation coefficients are labelled. **G**, Box plots showing the exon usage in every 15 cells ranking with cell areas from smallest to largest. Each dot represents a cell, colour is coded by the median usage in each bin. Box plots’ horizontal lines indicate the first, second (median) and third quartiles. **H**, Scatter plot of the maximum (y-axis) and minimum (x-axis) correlation coefficients between exon usage and cell area within a gene. Genes that are expressed with less than 10 cells are filtered out. Gene DAAM1 in (G) is labelled as red. Genes with smallest minimum correlation coefficient and largest maximum correlation coefficient are labelled with orange circles.

To investigate the relationship between cell size and transcriptional profile, the expression of each gene was correlated with cell size. Since our previous data suggested MT-high cells are apoptotic, these analyses used only MT-low cells. The distribution of correlation coefficients (Spearman ρ) peaked around zero, indicating that most genes are evenly expressed across cell sizes (**Figure 3B** and **Supplementary Figure 3B**). Some genes were expressed at lower levels in larger cells (e.g. RBM8A) (**Figure 3C**, left) and some genes at higher levels in larger cells (e.g. HIGD1A) (**Figure 3C**). To understand the biology of genes associated with cell size, the most enriched pathways within genes with largest positive or negative correlation with cell size were computed via GSEApy ^40^ (**Figure 3D**). Pathways associated with positive correlation included microtubule nucleation and polymerization ^42^, an indication that larger cells might be growing their cytoskeleton, perhaps in preparation for mitosis. Among the hits in this pathway was TUBG2, which encodes a protein required for microtubule nucleation at the centrosome (**Supplementary Figure 3C**). Two of the pathways associated with negative correlation related to RNA degradation. Genes in these pathways included RBM8A (**Figure 3C**, left), also known as Y14, and EIF4A3, both of which encode core members of the exon splicing junction complex (EJC) ^43,44^. EIF4A3 encodes a DEAD-box protein that is bound to ATP, while the MAGOH-Y14 heterodimer inhibits EIF4A3 ATPase activity, destabilising the association of the EJC with its RNA substrate.

Given that spliceosome factors were downregulated in larger cells, we suspected that cells of different size might show preferential usage of specific gene isoforms. To explore this hypothesis, HTSeq 2.0 was employed to subassign reads for individual genes into their exonic components ^45^. Within each gene, the fraction of reads assigned to each exon were computed in every cell (**Figure 3E**). Fractional exon usage was then correlated with cell size, yielding a zero-centred distribution (**Figure 3F**). To understand the meaning of this result, we considered DAAM1, a member of microfilament-related formins ^46^. While exon 11 showed a similar fractional usage across all cell sizes (**Figure 3G**, middle), exon 25 was preferentially used by smaller cells (**Figure 3G**, left) and exon 1 by larger ones (**Figure 3G**, right). The minimum and maximum correlation coefficients between exon usage and cell area were computed for each gene (**Figure 3H**). The majority of genes were located close to the origin, indicating little exonic preference across cell sizes. A streak of genes was close to the diagonal, which is consistent with a maximum-parsimony model with only two preferential exons (one for small cells, one for large cells). Some genes were located off-origin and off-diagonal, suggesting more complex splicing preferences. Across the entire transcriptome, origin recognition gene ORC1 showed the smallest minimum correlation, while long non-coding RNA KTN1-AS1 showed the largest maximum correlation.

In addition to cell size, cell eccentricity (**Supplementary Figure 3D**) was also computed and correlated with gene expression to exemplify the usage of more complex morphological features, resulting in a separate set of genes and pathways (**Supplementary Figures 3E-G**). Overall, these findings indicate that the MCF-7 cell size heterogeneity is associated with preferential usage of biological pathways, single genes, and individual exons.

### HyperSeq peers into the metabolic state of cells

The hyperspectral information from each cell can also be used to study specific autofluorescent biomolecules. Nicotinamide adenine dinucleotide (NADH) and its phosphate (NADPH), two enzyme cofactors and electron carriers playing ubiquitous roles in cell metabolism ^47,48^, are among the brightest autofluorescent biomolecules within human cells. A hyperspectral channel was therefore chosen to study NAD(P)H metabolism, aiming to correlate its intensity with gene expression from the same cell (**Figure 4A**). Because autofluorescence is an optically incoherent process involving vibrational quantum states of the medium, the signal detected in the microscope camera is proportional to the direct product of excitation spectrum times emission spectrum at the chosen pair of wavelengths (**Figure 4B**). Therefore, the hyperspectrum of free NAD(P)H was reconstructed from previously published spectra ^49^ (**Supplementary Figure 4A**) and intensity in channel 4 (*λ*_ex_ = 343 nm, *λ*_em_ = 451 nm) was chosen as a proxy for the spectral peak of this molecule (**Figure 4C**, left). The correlation between channel 4 intensity with gene expression resulted in a distribution of correlation coefficients with a slight positive bias (Spearman ρ, **Figure 4D**). Multiple genes related to the production of NADH had strong positive correlations with channel 4 intensity, including IDH1 (ρ=0.26) and ME2 (ρ=0.29) (**Figure 4E**). LDHA, which catalyses NADH conversion into NAD+, had a strong negative correlation (ρ=-0.31, **Figure 4E**). These data indicate that hyperspectral imaging in channel 4, which is proximal to the NAD(P)H autofluorescence spectral peak, is consistent with a specific transcriptomic signature affecting NAD(P)H metabolism. We also performed a separate analysis on the ratio between channel 4 and channel 2 intensities, which is expected to contain information about the blue-shifted spectrum of protein bound NAD(P)H ^50,51^ (**Figure 4C**, right). That analysis indicated a more narrow variation in correlation coefficient and a partially overlapping set of genes (**Supplementary Figures 4B-D**). MCF-7 cells utilise glucose for about half of their oxidative metabolic energy needs, which requires TCA cycle enzymes to generate mitochondrial NADH ^52^. Therefore, we examined the distribution of correlation coefficient in TCA cycle and nucleotide synthesis showed a slight tendency towards lower correlation coefficients (**Figure 4F, Supplementary Table 2**). Taken together, these data suggest that HyperSeq can be used to inform about the biology of genes and pathways directly and indirectly related to a chosen autofluorescent metabolite.

**Figure 4:**
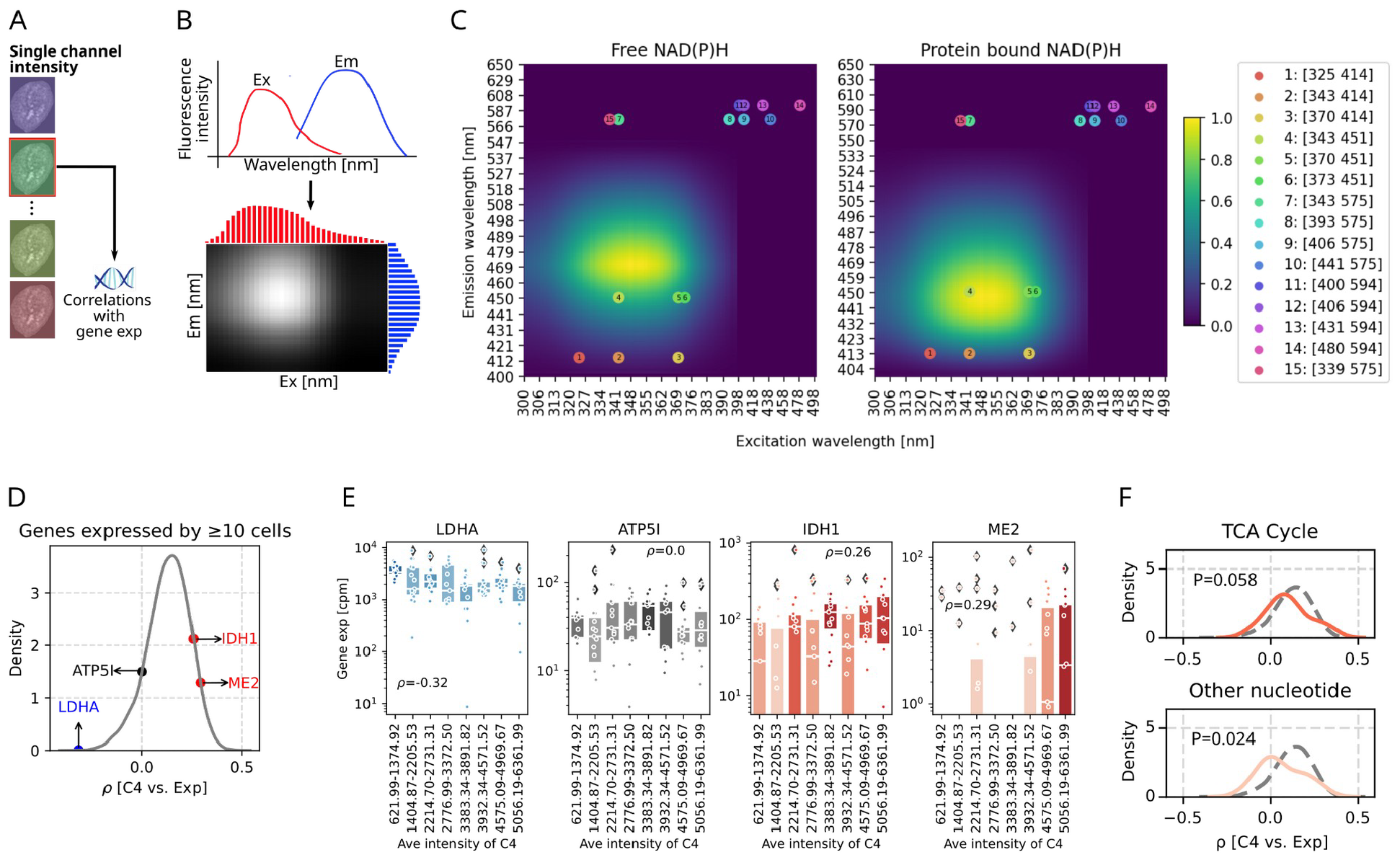
HyperSeq generates hypotheses about cell metabolism. **A**, Schematic of workflow to correlate a single channel intensity with gene expression. **B**, Schematic plot of constructing spectra matrix using the excitation and emission spectra. **C**, Spectra matrices of free (left) and protein bound NAD(P)H (right). Dots represent the positions of the 15 hyperspectral channels according to the excitation and emission wavelengths, and numbers inside the dots correspond with the channel number in the legend box. **D**, KDE plot of correlation coefficients between spectra intensities in channel 4 and gene expressions for genes expressed by ≥ 10 cells. Four genes with negative, ∼ zero, and positive correlation coefficients are labelled. **E**, Box plots showing the gene expression (cpm) in every 15 cells ranking with spectra intensities in channel 4 from smallest to largest of genes in (D). Each dot represents a cell, and box colour is coded by the median expression in each bin. Box plots’ horizontal lines indicate the first, second (median) and third quartiles. **F**, KDE plots showing the correlation coefficients between spectra intensity in channel 4 and gene expression of genes involved in two cellular metabolic pathways. Vertical grey dashed lines indicate correlation coefficient of - 0.5, 0, 0.5, while the horizontal grey dashed line indicates a KDE density of 5. The grey dashed curve presents the KDE distribution for all genes in the pathway (as in D). The colour of the KDE distribution is coded by the correlation coefficient of the peak.

## Discussion

The integration of omics data with complementary measurements from the same cell has been limited by challenges in combining high-throughput single-cell assays into a single experimental approach ^53,54^. Spatial transcriptomic technologies ^27,55^, while growing in popularity, currently suffer from either low spatial resolution or low RNA capture efficiency. Live-seq, which uses an atomic force microscope ^13^, can be time-consuming and requires a high level of expertise. Chip-based techniques are addressable and high-throughput but have limitations in imaging resolution ^33,56^. HyperSeq was designed to make integration of imaging and sequencing accessible to a broader audience of biomedical researchers. Notably, HyperSeq separates the high-resolution imaging platform from the cell picker, which lifts the requirement to have a single, highly specialised setup. In our case, the two pieces of equipment were on adjacent floors of one building. It might be possible to transport the cells further if media is added after imaging. DAPI can be used to increase the fraction of high-quality cells captured (**Supplementary Figure 1D**) to almost 100%, similar to fluorescence activated cell sorting (FACS) ^57^. Future research is planned to extend the current scope to other cell lines, cells in suspension, mixed cell populations, and primary cells. Other potential extensions of HyperSeq include chromatin accessibility ^12,58^, whole-genome sequencing ^59^ at the sequencing level, and analysis of subcellular organelles ^60,61^ at imaging level. Hyperseq could also be used on engineered cells that export samples of the transcriptome nondestructively ^62^, potentially achieving data sets similar to Live-seq with a simpler workflow.

HyperSeq was used to explore transcriptomic differences across hyperspectral heterogeneity. Current applications of hyperspectral imaging remain at the stage of discerning different physiological states of cells and tissues based on their autofluorescent hyperspectrum, for example the assessment of the viability of islets ^63^ and watermelon seeds ^64^. However, research aimed at probing the molecular distinctions underlying specific hyperspectral patterns is limited. In this study, we identified a subset of cells displaying multiband autofluorescence rings at the plasma membrane that also presented an altered transcriptome, with potentially upregulation of gene silencing and down regulation of cellular respiration. This result provides a blueprint for future studies using HyperSeq to reveal the transcriptomic changes underpinning spectral changes in specific subcellular compartments, such as the nucleus or mitochondria.

HyperSeq can also be used to correlate gene expression with brightfield images. Regulation of cell morphology is complex ^65,66^, therefore technical advances in this area would be important to better our understanding. HyperSeq is also applicable to investigate the relationship between alternative splicing and visual phenotypes such as cell morphology or autofluorescence. Unlike droplet-based methods ^10,7^ which generate reads at one end of the transcript, HyperSeq reads cover the entire gene body ^67^, enabling read counting on a per-exon basis using HTSeq 2.0 ^45^ or similar tools. Although other computational approaches to single cell isoform analysis such as SingleSplice ^68^, BRIE ^69^, and Expedition ^70^ were not assessed in this study, they are expected to be compatible with HyperSeq because the genomics data is in the same format as Smart-seq 2 ^57^. More tentatively, it might be possible to adapt the library preparation protocol to long-read sequencing technologies (e.g. PacBio, Oxford Nanopore) ^67^ to perform association analysis between full isoform abundance and hyperspectral features.

The transcriptomic proxies in living cells of autofluorescent biomolecules such as NAD(P)H can also be studied using HyperSeq. NAD(P)H plays ubiquitous roles in cell metabolism ^47,48^. In addition, NADH binding sites are known to be altered by metabolic pathways related to carcinogenesis and differentiation, and enzymatic binding directly influences NADH cycling through energy production pathways ^71^. For example, Bird et al. found that in breast cancer cells, the ratio of free to protein-bound NADH is related to the NADH/NAD+ redox ratio ^72^. In this study, we peered into the metabolic states of MCF-7 cells by correlating transcriptomic expression with features extracted on hyperspectral channels designed for NAD(P)H. While this study has focused on NAD(P)H, other biomolecules of importance for cell metabolism and behaviour could be studied using the same concept, including flavins, collagen and protoporphyrin IX ^73,74^. A limitation of this study is that optical features were not background-subtracted and calibrated into absolute metabolite concentrations. Future work is planned to address this point.

In conclusion, HyperSeq was shown to be a robust approach to integrate hyperspectral imaging and transcriptomics at the single cell level, illuminating the hidden relationship between a cell’s appearance and its internal molecular state.

## Acknowledgements

We would like to thank Tatyana Chtanova for scientific discussions. This work was supported by a UNSW Cellular Genomics Futures Institutes Seed Grant (SF-014) to F.Z.. Y.X. was supported by an Australian Government Research Training Program (RTP) Scholarship.

## Methods

### Cell culture

The human breast cancer cell line MCF-7 was purchased from American Type Culture Collection. MCF-7 cells were cultured in RPMI (Gibco, Carlsbad CA, USA) supplemented with 10% fetal bovine serum (FBS, Bovogen, Sydney, Australia) and 4mM L-glutamine (Thermo Fisher Scientific, Waltham MA, USA). The cells were seeded in a 35 mm coverslip-bottomed dish with an imprinted 500μm cell location grid (ibidi, Gräfelfing, Germany) and cultured at 37°C in a humidified atmosphere with 5% CO_2_ to adhere overnight. To avoid overcrowding, which can increase the rate of doublet or multiplet capture due to strong cell-cell adhesion, cell density was kept at ∼ 5,000 cells per dish (**Supplementary Figure 1D**).

### Hyperspectral imaging

MCF-7 cells were washed twice with PBS, then placed into equilibrated Hank’s balanced salt solution (HBSS) (Thermo Fisher Scientific, Waltham MA, USA) and maintained at 37°C during imaging. Hyperspectral imaging was carried out with a customised fluorescence microscope (Olympus IX83, Southall, UK) with a 40× oil objective (NA 1.35, Olympus, Southall, UK), an iXon Ultra 888 EMCCD (model: hnu1024, Andor, Perthshire, Scotland) and operated below −30°C to reduce noise. Custom-designed LED illumination and filter cubes enabled autofluorescence imaging in 15 channels (**Supplementary Table 1**; excitation values are ± 5nm and emissions are ± 20 nm).

### Single cell capture

To capture cells after imaging, both manual and automatic modes of CellCelector (Sartorius AG, Niedersachsen, Germany) were used. After hyperspectral imaging, MCF-7 cells in the gridded dish were washed twice with PBS. To reduce adherence of MCF-7 cells without causing excessive motility, cells were treated with 10% TrypLE™ Express Enzyme (no phenol red, Thermo Fisher Scientific, Waltham MA, USA) for 30 min at room temperature before capture.

Preliminary tests indicated that specific concentration and treatment time might be required for different cell lines. Cells were identified on the CellCelector platform, picked by the 30μm glass capillary under manual picking mode. The captured cells were dispensed with 0.2μL DNase/RNase-Free Water (Thermo Fisher Scientific, Waltham MA, USA) into a 384-well Frame-Star plate (4titude Ltd, Surrey, UK) containing 1μL lysis buffer. The lysis plate was cooled down to 4ºC at the rack tray during cell capture, and was stored at −80ºC for subsequent procedures. In batch 2, DAPI (1:1000) (Thermo Fisher Scientific, Waltham MA, USA) was added into PBS supplemented with 10% TrypLE™ Express Enzyme 10 min before cell capture. Both DAPI and brightfield channels of CellCelector were used to identify cells. Inside the imaged grids, cells without DAPI fluorescence were captured into a 384-well Frame-Star plate with 1μL lysis buffer. All the other procedures are the same.

### Single cell transcriptomic library preparation

Library was prepared following Smart-seq2 protocol ^57^. Briefly, cells were picked into 384 well lysis plates followed by reverse transcription, template switching and 26 cycles of PCR to generate and amplify the cDNA. MANTIS automated liquid handler (Tecan, Zürich, Switzerland) was used during the process. cDNA was normalised to 0.1 - 1ng/μl according to the qualification by Quant-iT™ PicoGreen™ dsDNA Assay Kit (Thermo Fisher Scientific, Waltham MA, USA) via automated liquid handling robot Mosquito (SPT Labtech, Melbourn, Cambridgeshire, UK). Nextera XT kit (Illumina, San Diego, CA, USA) with 14 cycles of amplification was used for tagmenting MCF-7 cells. Purification was conducted to remove primer by Agencourt Ampure XP (Beckman Coulter, Brea CA, USA) magnetic beads at a ratio of 0.7x for two times. Libraries were quantified using Bioanalyzer 2100 (Agilent Technologies, Waldbronn, Germany).

### Next-generation sequencing

All libraries were sequenced at Ramaciotti Centre for Genomics (UNSW). Library of batch 1 was sequenced on NextSeq 500 (Illumina, San Diego, CA, USA) using 75 base paired-end reads. Library of batch 2 was sequenced on NextSeq 1000 (Illumina, San Diego, CA, USA) using 150 base paired-end reads. Library of the pilot experiment was sequenced on MiSeq (Illumina, San Diego, CA, USA) using 75 base paired-end reads. Sequencing coverage was around 500,000 to 5,000,000 read pairs per cell.

### scRNA-seq data analysis

The following open source software was used for this study: numpy ^75^, pandas ^76^, matplotlib ^77^, seaborn ^78^, scipy ^79^ and scanpy ^36^.

### Pre-processing of scRNA-seq data

The sequencing reads were demultiplexed using bcl2fastq 2.20 (Illumina, San Diego, CA, USA), mapped and counted to human genome reference (GRCh38) for MCF-7 cells using STAR aligner ^80^. HTSeq 2.0 ^45^ was used to map the sequencing reads of MCF-7 cells into exon level. Quality of datasets were evaluated based on the number of genes expressed in the count matrix, the total counts per cell and the percentage of counts in mitochondrial genes. Genes that were expressed in less than ten cells were excluded. Datasets were normalised by the total reads, multiplied by 1,000,000 (cpm). GSEAPY ^40^ was used for pathway analysis.

### Image analysis and cell segmentation

Hyperspectral figures were normalised by the most frequent fluorescence intensity in each channel, which was regarded as the intensity of background, and then multiplied by 30, a close intensity as the background. Pseudo RGB figures were created using channels 1, 6, 13 to clarify cells from the background. Rough positions of single cells were obtained via the Pixel Classification function in Ilastik ^81^ (**Supplementary Figure 1C, left**) based on these fake RGB figures. A customised semi-automatic GUI written based on tkinter ^82^ was designed to define the cell boundary (red dots in **Supplementary Figure 1C, right**).

### Validation of cells inside the lower-left subgroup

To distinguish between the two cell subpopulations identified by PC1 and PC2 from PCA analysis of the 15 average optical features of all cells, the boundary of the lower-left subgroup was determined as intensity value of 0.18 from kernel density estimation (KDE) analysis using scipy ^79^ (**Supplementary Figure 1E**).

### Validation of cells with bright border rings

To determine whether a cell exhibits a bright border ring at the plasma membrane, the border ring and inner ring adjacent to the border ring, each with a thickness of 10 pixels (**Supplementary Figure 2B**), were segmented. Subsequently, log2 fold changes between the average fluorescence intensity of the border and inner rings in channel 13 (*λ*_ex_ = 431nm, *λ*_em_ = 594nm) were computed. Cells with log2 fold changes greater than 0.4 were categorised as ‘ringed cells’, while those with log2 fold changes less than 0.4 were categorised as ‘unringed cells’.

### Hyperspectral imaging feature extraction

#### Machine learning features

Principal component analysis ^37^ was conducted on 15 average optical features of all cells using scipy ^79^. Computational features with high cumulative explained variances were selected for further analysis.

#### Morphological features

Cell area and cell eccentricity that describe cell morphology were designed. Cell area was computed as the pixel counts of the segmented cell (**Figure 3A**), while eccentricity that reflects cell shape was calculated as y/x-1 (y: cell length, x: width) (**Supplementary Figure 3D**).

#### Optical features

Images of segmented cells were first processed by Gaussian blur in scikit-image ^83^ to reduce noise. Average fluorescence intensities of each cell in each channel were computed as optical features (**Figure 4A**). Intensity ratios were defined as the ratios between average fluorescence intensities of two different channels (**Supplementary Figure 4B**).

### Construction of spectra matrices of autofluorescent biomolecules

In order to get the absorbance of molecules in multiple channels, the excitation (Ex) and emission (Em) spectrum of molecules were downloaded from published papers ^49^. Spectra matrices were constructed based on both Ex and Em spectrums (**Figure 4B**). In particular, the fluorescent intensities at different Ex and Em wavelengths are calculated according to Equation 1,

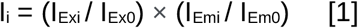

where Ex0 and Em0 represent the Ex and Em with the highest fluorescent intensity in the spectrum, respectively, Exi and Emi are the Ex and Em under the ith condition, I_i_: normalised fluorescence intensity at Exi and Emi, I_Exi_ and I_Ex0_ represent the fluorescent intensity of Exi and Ex0 in the Ex spectrum, and I_Emi_ and I_Em0_ mean the fluorescent intensity of Emi and Em0 in the Em spectrum.

### Correlation analyses between hyperspectral, morphological, and optical features and gene expression/exon usage

In order to keep enough genes for further study, genes expressed by ≥10 cells were selected for correlation analyses. The expression of each gene was correlated with distinct features by Spearman’s correlation in MT-low cells only (117 cells in total). The distribution of correlation coefficients became closer to a normal distribution with an increase in the number of cells expressing genes because of reduced noise from ties in the cell ranks. Box plots were created to display representative genes from correlation analyses with gene expression (cpm) in cells, ranked from smallest to largest based on various features. The first seven boxes each comprised 15 cells, while the last box contained 12 cells.

## Data and Code Availability

Python 3 and Jupyter notebooks were used for the analysis and are available at https://github.com/echosun77/MCF7_hyperspectral_imaging_sequencing. The sequencing reads and the full data set including images and count matrices were deposited on GEO with Series id GSE254034.

