## Supplementary Figures for "Integration of hyperspectral imaging and transcriptomics from individual cells with HyperSeq"

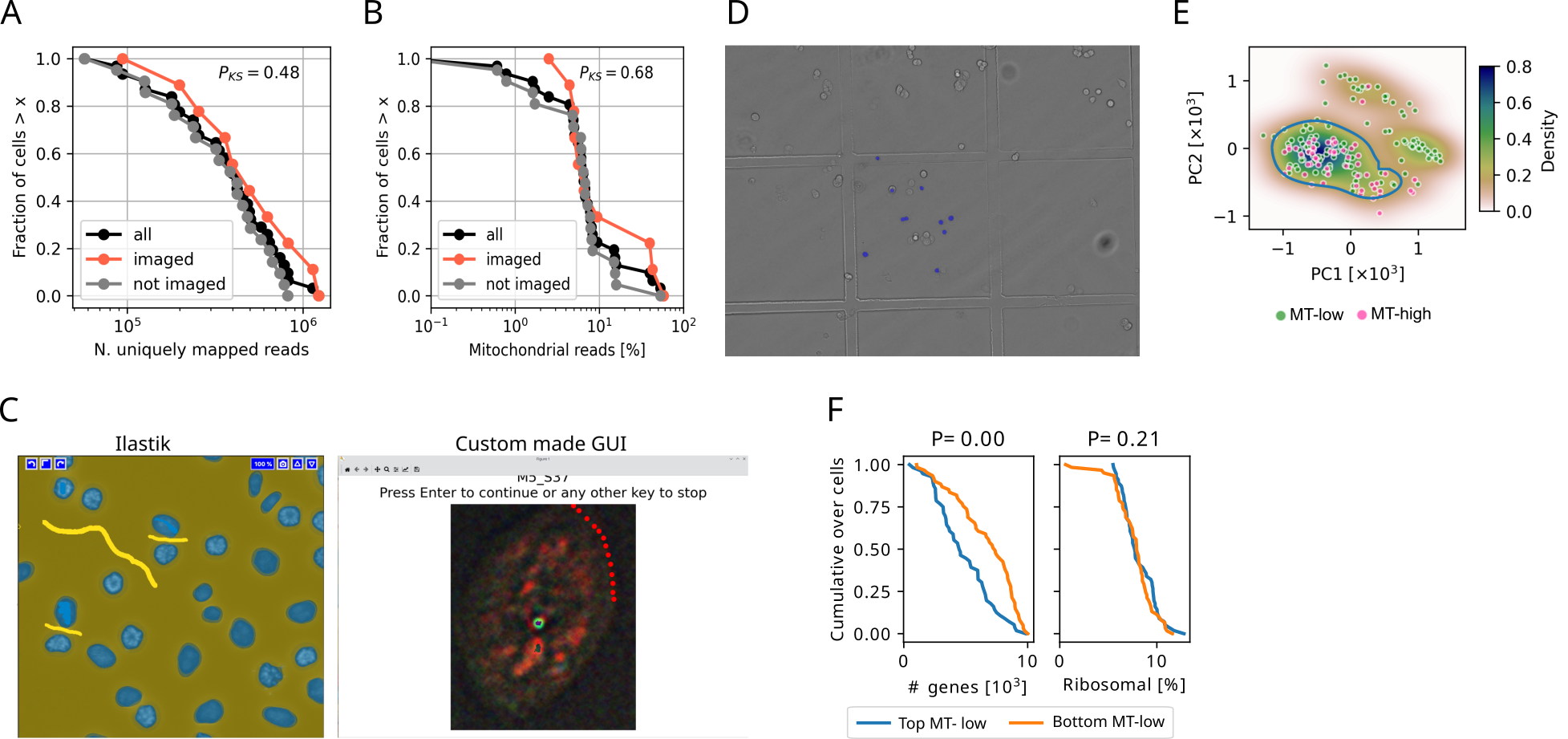
**Supplementary Figure 1.** **A, B**, RNA quality control by comparing imaged cells and non-imaged cells in the pilot experiment. **C**, Schematic plots of Ilastik (left) and the custom made GUI (right) for cell segmentation. **D**, Image of MCF-7 cells on the gridded plate after imaging. This image was constructed by both brightfield and DAPI fluorescent images. Cells masked with blue colour indicate cells with DAPI signal under fluorescent microscope. The DAPI mask was defined by the threshold of DAPI fluorescent intensity. **E**, Scatter plot of PC1 and PC2 of each cell. The contour of the lower-left subgroup was drawn as the density from KDE analysis on PC1 and PC2 as 0.18. **F**, Distribution of total counts, the number of genes, and percentage of MT-low cells inside or outside of the lower-left subgroup.

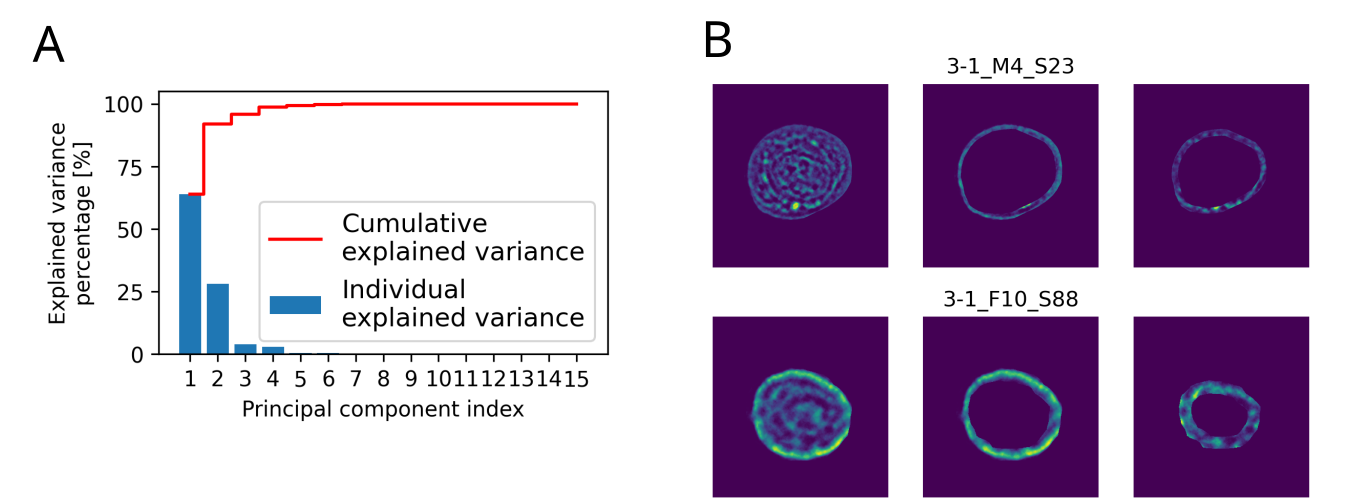
**Supplementary Figure 2.** **A**, Barplot and cumulative plot showing the explained variance percentage (%) of each principal component (PC) from PCA on 15 channels. **B**, Schematic plot of the border and inner rings segmented by distance of 10 pixels of cells without (the cell in **Figure 2B**, upper panel) and with (the cell in **Figure 2B**, lower panel) bright rings in channel 13.

**Supplementary Figure 3.** **A**, Figure showing areas of all cells based on the number of pixels from lowest to largest. **B**, KDE plot of correlation coefficients between cell area and gene expression on genes expressed by ≥ 0, 10, and 50 cells. **C**, Box plot showing the expression (cpm) of TUBG2 in every 15 cells ranking with cell areas from smallest to largest of a gene exclusively expressed in larger cells. Each dot represents a cell, and box colour is coded by the median expression in each bin. Box plots’ horizontal lines indicate the first, second (median) and third quartiles. **D**, Schematic plot of computing cell eccentricities. **E**, KDE plot of correlation coefficients between cell eccentricity and gene expression on genes expressed by ≥ 0, 10, and 50 cells. **F**, KDE plot of correlation coefficients between cell eccentricity and gene expression for genes expressed by ≥ 10 cells. Three genes with negative, ~zero, and positive correlation coefficients are labelled. **G**
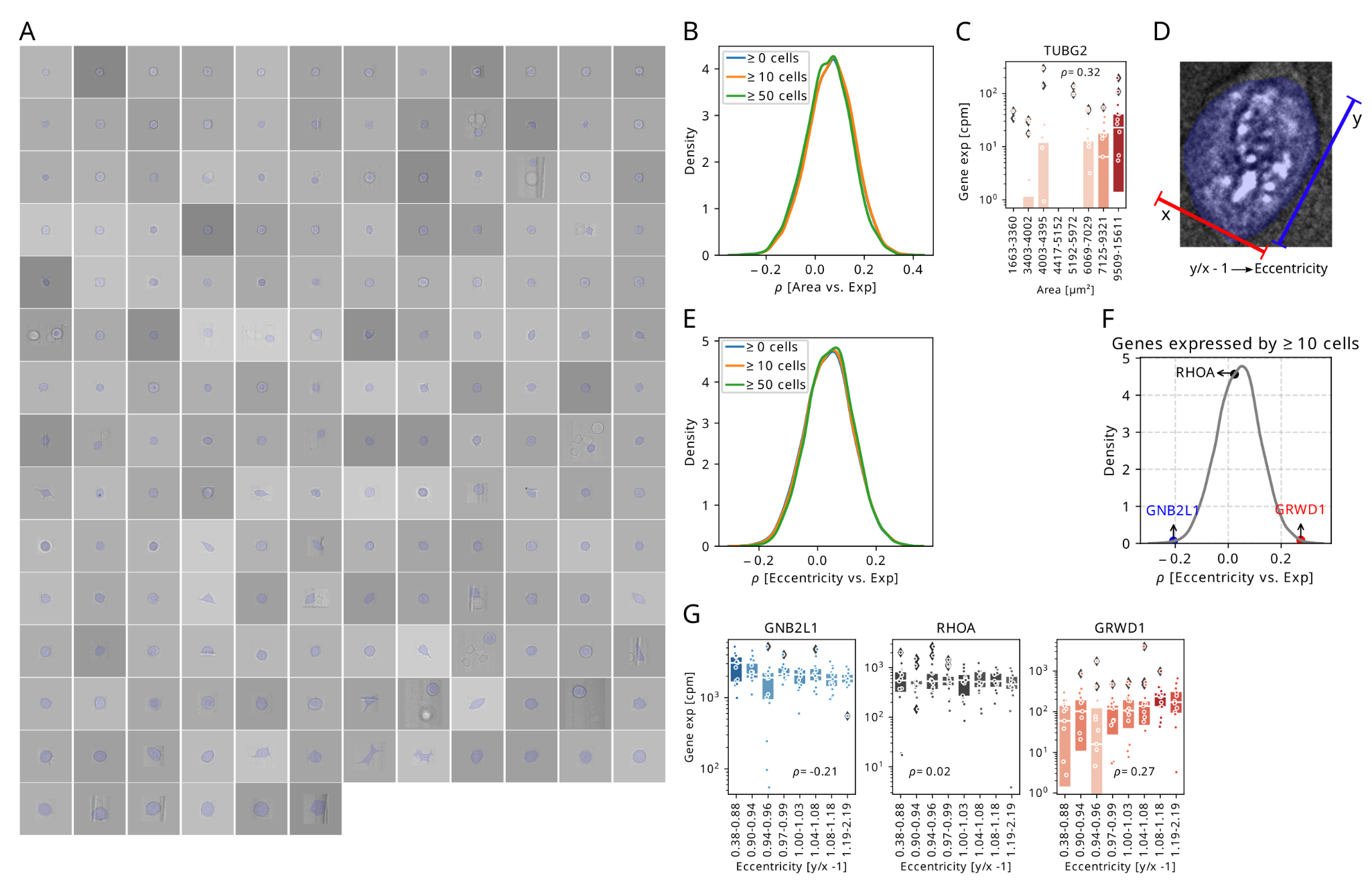
, Box plots showing the gene expression (cpm) in every 15 cells ranking with cell eccentricities from smallest to largest of genes in (F). Each dot represents a cell, and box colour is coded by the median expression in each bin. Box plots’ horizontal lines indicate the first, second (median) and third quartiles.

**Supplementary Figure 4. A**, The absorbance (left) and respective (right) emission spectra of typical autofluorescent molecules with NADH highlighted. The height of the peaks indicates the relative molar absorption coefficient and the quantum yield, respectively. Image from ^49^. **B**, Schematic plots of correlations between the intensity ratio and gene expression. **C**, KDE plot of correlation coefficients between the ratio of spectra intensities in channel 4 to channel 2 and gene expressions for genes expressed by ≥ 10 cells. Three genes with negative, ~ zero, and positive correlation coefficients are labelled. **D**
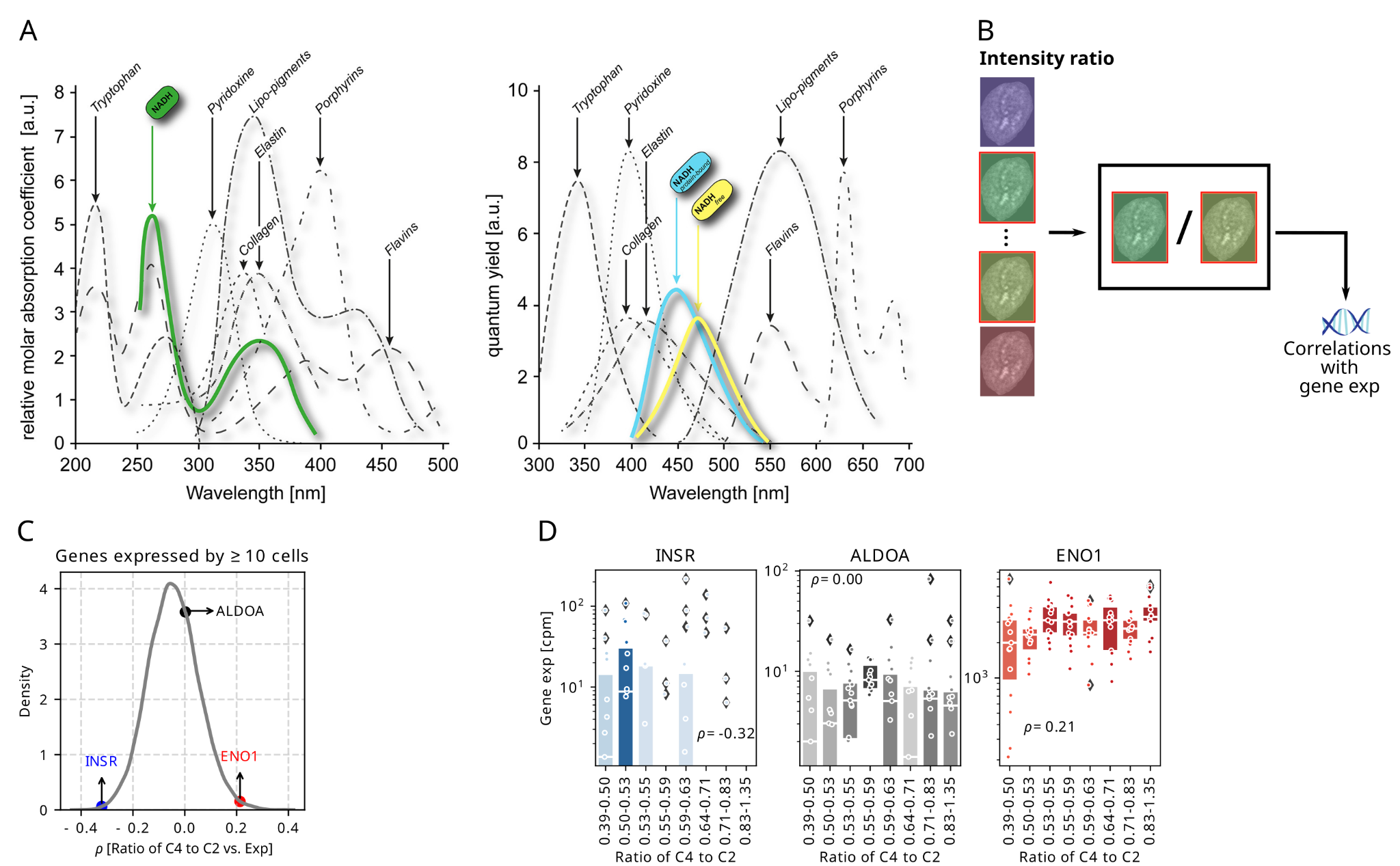
, Box plots showing the gene expression (cpm) in every 15 cells ranking with the ratio of spectra intensities in channel 4 to channel 2 from smallest to largest of genes in (C). Each dot represents a cell, and box colour is coded by the median expression in each bin. Box plots’ horizontal lines indicate the first, second (median) and third quartiles.
