## Supplementary Tables for "Integration of hyperspectral imaging and transcriptomics from individual cells with HyperSeq"

**Supplementary Table 1: Specification of spectral channels with their respective excitation, emission, and exposure time for the hyperspectral system.**

| Channel | Central wavelength (nm) | | Exposure time (s) |
| --- | --- | --- | --- |
|  | Excitation | Emission |  |
| 1 | 325 | 414 | 0.5 |
| 2 | 343 | 414 | 0.2 |
| 3 | 370 | 414 | 0.1 |
| 4 | 343 | 451 | 0.05 |
| 5 | 370 | 451 | 0.01 |
| 6 | 373 | 451 | 0.01 |
| 7 | 343 | 575 | 0.2 |
| 8 | 393 | 575 | 0.01 |
| 9 | 406 | 575 | 0.02 |
| 10 | 441 | 575 | 0.03 |
| 11 | 400 | 594 | 0.05 |
| 12 | 406 | 594 | 0.02 |
| 13 | 431 | 594 | 0.02 |
| 14 | 480 | 594 | 0.05 |
| 15 | 339 | 575 | 0.001 |

**Supplementary Table 2: Curated gene lists in the TCA cycle and nucleotide synthesis pathways.**

| **TCA Cycle** | **Other nucleotide synthesis** |
| --- | --- |
| IDH2 | NME1 |
| MDH2 | CMPK1 |
| SDHA | UPP1 |
| CS | RRM2 |
| MPC1 | UCK2 |
| SDHC | UCKL1 |
| SUCLG1 | TK1 |
| ME1 | RRM1 |
| SDHB | UCK1 |
| IDH3G | UNG |
| IDH3B | RRM2B |
| MPC2 |  |
| ACO2 |  |
| MDH1 |  |
| DLD |  |
| OGDH |  |
| SDHD |  |
| PDHB |  |
| DLAT |  |
| ACO1 |  |
| FH |  |
| ACLY |  |
| PC |  |
| SUCLA2 |  |
| IDH3A |  |
| IDH1 |  |
| DLST |  |
| ME2 |  |
| PDHA1 |  |
| SUCLG2 |  |
